# A systematic review of Hepatitis B virus (HBV) drug and vaccine escape mutations in Africa: a call for urgent action

**DOI:** 10.1101/258350

**Authors:** Jolynne Mokaya, Anna L McNaughton, Martin J Hadley, Apostolos Beloukas, Anna-Maria Geretti, Dominique Goedhals, Philippa C Matthews

**Affiliations:** Nuffield Department of Medicine, University of Oxford, Medawar Building, South Parks Road, Oxford OX1 3SY, UK; Oxford University Academic IT Department, 13 Banbury Road, Oxford OX2 6NN, UK; Institute of Infection and Global Health, 8 West Derby Street, University of Liverpool, Liverpool L69 7BE, UK; Division of Virology, University of the Free State/National Health Laboratory Service, PO Box 339(G23), Bloemfontein 9300, Republic of South Africa; Department of Microbiology and Infectious Diseases, Oxford University Hospitals NHS Foundation Trust, John Radcliffe Hospital, Headley Way, Headington, Oxford OX3 9DU, UK

**Keywords:** Hepatitis B virus, treatment, antiviral, nucleotide analogues, mutations, drug resistance, vaccine escape, Africa

## Abstract

International sustainable development goals for the elimination of viral hepatitis as a public health problem by 2030 highlight the pressing need to optimize strategies for prevention, diagnosis and treatment. Selected or transmitted resistance associated mutations (RAMs) and vaccine escape mutations (VEMs) in hepatitis B virus (HBV) may reduce the success of existing treatment and prevention strategies. These issues are particularly pertinent for many settings in Africa where there is high HBV prevalence and co-endemic HIV infection, but lack of robust epidemiological data and limited education, diagnostics and clinical care. The prevalence, distribution and impact of RAMs and VEMs in these populations are neglected in the current literature. We therefore set out to assimilate data for sub-Saharan Africa through a systematic literature review and analysis of published sequence data, and present these in an on-line database (https://livedataoxford.shinyapps.io/1510659619-3Xkoe2NKkKJ7Drg/). The majority of the data were from HIV/HBV coinfected cohorts. The commonest RAM was rtM204I/V, either alone or in combination with compensatory mutations, and identified in both reportedly treatment-naïve and treatment-experienced adults. We also identified the suite of mutations rtM204V/I + rtL180M + rtV173L, that has been associated with vaccine escape, in over 1/3 of cohorts. Although tenofovir has a high genetic barrier to resistance, it is of concern that emerging data suggest polymorphisms that may be associated with resistance, although the precise clinical impact of these is unknown. Overall, there is an urgent need for improved diagnostic screening, enhanced laboratory assessment of HBV before and during therapy, and sustained roll out of tenofovir in preference to lamivudine alone. Further data are needed in order to inform population and individual approaches to HBV diagnosis, monitoring and therapy in these highly vulnerable settings.

**Author’s summary:** The Global Hepatitis Health Sector Strategy is aiming for the elimination of viral hepatitis as a public health threat by 2030. However, mutations associated with drug resistance and vaccine escape may reduce the success of existing treatment and prevention strategies. In the current literature, the prevalence, distribution and impact of hepatitis B virus (HBV) mutations in many settings in Africa are neglected, despite the high prevalence of HBV and co-endemic HIV infection. This systematic review describes the frequency, prevalence and co-occurrence of mutations associated with HBV drug resistance and vaccine escape mutations in Africa. The findings suggest a high prevalence of these mutations in some populations in sub-Saharan Africa. Scarce resources have contributed to the lack of HBV diagnostic screening, inconsistent supply of drugs, and poor access to clinical monitoring, all of which contribute to drug and vaccine resistance. Sustainable long-term investment is required to expand consistent drug and vaccine supply, to provide screening to diagnose infection and to detect drug resistance, and to provide appropriate targeted clinical monitoring for treated patients.

## INTRODUCTION

In 2015, the World Health Organisation (WHO) estimated that 3.5% of the world’s population (257 million people) were living with Hepatitis B virus (HBV) infection, resulting in 887,000 deaths each year, mostly from complications including cirrhosis and hepatocellular carcinoma (HCC) [1]. United Nations Sustainable Development Goals set out the challenge of elimination of viral hepatitis as a public health threat by the year 2030 [2]. One of the existing strategies in the elimination toolbox is use of antiviral drugs in the form of nucleos(t)ide analogues (NAs). Suppression of viraemia not only reduces inflammatory and fibrotic liver disease in the individual receiving treatment but also reduces the risk of transmission. However, the emergence of HBV resistance-associated mutations (RAMs) is a potentially significant concern for the success of this strategy.

Africa is the continent with the second largest number of individuals with chronic HBV (CHB) infection, with an estimated 6.1% of the adult population infected [1]. However, there is little commitment and resource invested into the burden of this disease, and many barriers are contributing to the epidemic [3,4]. Globally, less than 10% of the population with CHB are diagnosed, with an even smaller proportion on treatment [1,4]. This proportion is likely to be even lower in Africa. The situation in Africa is further complicated by the substantial public health challenge of coendemic human immunodeficiency virus (HIV) and HBV; coinfection worsens the prognosis in dually infected individuals [5]. There is also a lack of robust epidemiological data on HBV from Africa [3,4].

Widespread use of antiretroviral therapy (ART) for HIV, incorporating NAs that also have activity against HBV, may have an impact on HBV through improved rates of viraemic suppression, but also potentially by driving the selection of RAMs. The WHO recommends screening for Hepatitis B virus surface antigen (HBsAg) in all HIV-1 infected individuals prior to ART initiation, and for all pregnant women during antenatal visits, to improve the clinical outcomes of people living with CHB and to enhance interventions that reduce the incidence of new cases [6]. However, screening of HBsAg is not routinely performed in many settings in Africa, with lack of implementation at least partially driven by cost and lack of programmes for HBV treatment outside the setting of HIV coinfection. HBV infected patients either remain untreated (most typical in the setting of monoinfection), or are exposed to antiviral drugs without proper monitoring and often intermittently, putting them at risk of developing RAMs (more likely in the setting of HIV coinfection) [4,7–10].

HBV is a DNA virus that replicates via an RNA intermediate, with reverse transcriptase (RT) catalysing the transcription of RNA into DNA [7]. NAs that inhibit RT are therefore used to prevent HBV replication, including lamivudine (3TC), entecavir (ETV) and tenofovir (conventionally in the form of tenofovir disoproxil fumarate (TDF), but more recently available as the prodrug, tenofovir alafenamide fumarate (TAF)), with mostly historical use of other agents including telbivudine (LdT) and adefovir (ADV) [6,11]. Choice of TDF/TAF or ETV is determined by availability, cost, safety profile and barrier to resistance [4]. In Africa, the choice of agent is usually limited to 3TC and TDF. Emergence of mutations happens because the RT enzyme is error-prone and lacks the proofreading function required to repair errors during transcription [7]; when these mutations confer a selective advantage by allowing the virus to escape the effect of drug therapy, they will become amplified in the viral population. Some RAMs confer resistance to one agent only, while others are associated with resistance to several agents (Fig 1).

**Fig 1:**
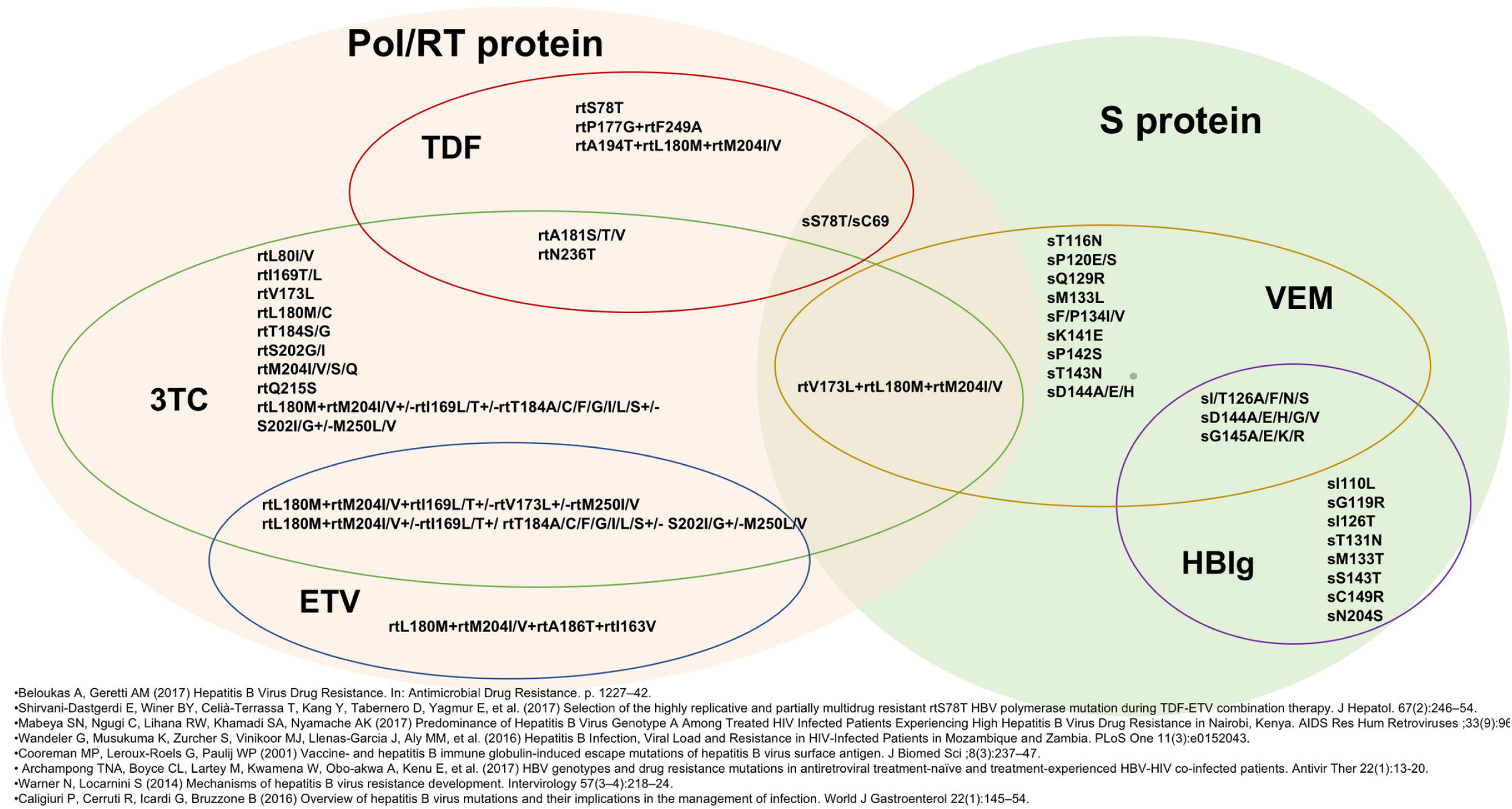
HBV drug resistance associated mutations (RAMs), vaccine escape mutations (VEMs) and mutations associated with Hepatitis B immunoglobulin (HBIg) resistance. HBV genes are shown in the coloured ovals. TDF = tenofovir, ETV = entecavir, 3TC = lamivudine. This figure incorporates data from eight studies; three were identified by the systematic review presented in this manuscript [12–14] and five from the wider literature [7, 15–18].

3TC was originally seen as a major breakthrough in treating HBV [19]. However, it is now known to have a low genetic barrier to resistance and its long-term effectiveness is limited as a result of resistance mutations in the ‘YMDD’ motif (tyrosine, methionine, aspartate, aspartate; amino acids 203-206) in domain C of the viral polymerase (Pol). These occur with associated upstream compensatory mutations in Pol domains A, B and in the B-C interdomain [7,15,16]. Among chronic HBV monoinfected patients, incidence of HBV resistance to 3TC is as high as 20% per year. In HIV/HBV coinfected patients, this can reach 90% over 5 years of treatment, as development of resistance is accelerated in HIV coinfection [5,20]. 3TC has also been associated with the induction of cross-resistance to emtricitabine (FTC), LdT, and at least partially ETV, thus reducing the options for subsequent treatment [10].

TDF is widely used in treatment of both HIV and HBV and is generally well tolerated. TDF has a high genetic barrier to resistance and maintains effective suppression of HBV in both monoinfected and HIV/HBV coinfected individuals [5,7,10,21,22]. Although it has a recognised association with nephrotoxicity in HIV treatment, current literature suggests it may be better tolerated in HBV infection [11]. Conversely, African populations have a higher background of renal disease [23] and could be potentially more vulnerable to nephrotoxitiy from TDF [24]. TAF delivers equally potent viraemic suppression at lower plasma levels, and is therefore associated with reduced nephrotoxicity [25], but is not available in Africa at present. HBV resistance to TDF is not well characterised, but there are emerging data from *in vitro* studies associating Pol mutations rtA194T and rtN236T with decreased susceptibility [11,21]. Virological breakthrough on TDF therapy has been reported in two patients harbouring rtS78T/sC69 mutations [17], and in another patient with multi-site polymerase mutations; rtL80M, rtL180M, rtM204V/I, rtA200V, rtF221Y, rtS223A, rtT184A/L, rtR153Q, and rtV191I [26]. The significance of these mutations needs to be further explored in clinical studies.

First line ART treatment regimens for HIV in sSA now almost universally include TDF, and current guidelines also recommend TDF-based regimens in individuals with HBV/HIV coinfection [27]. Accordingly, in both HIV monoinfection and HBV/HIV coinfection, use of TDF has increased across much of Africa. Nevertheless, it remains the case that 3TC is used as the only HBV-active agent in some settings [7,8], as well as in second line regimens, exemplified by South Africa where second line ART substitutes Zidovudine (AZT) for TDF leaving only 3TC coverage for HBV [28]. Among HBV/HIV coinfected children in South Africa treated with regimens including 3TC and/or TDF, HBV viraemia has been demonstrated, highlighting potential underlying HBV drug resistance [29].

ETV is another active agent, and is safe and well tolerated. However it is not active against HIV and therefore has to be added to ART regimens rather than being part of the primary backbone, is not recommended in pregnancy, and is not routinely available in most African settings [30]. Resistance arises more commonly in the context of prior 3TC exposure [11,31], which may limit its future potential in Africa, particularly in HIV endemic populations.

As a component of the Expanded Programme on Immunization (EPI), HBV preventive vaccines have been rolled out in Africa since 1995 [4]. HBV vaccine is highly effective in prevention of mother to child transmission (PMTCT); when administered to infants within 24 hours of birth followed by a dose given at 6 and another at 14 weeks to complete the primary series, it reduces the rate of mother to child transmission by 85% - 95% [32,33]. However, by 2016 only 11 countries in Africa had adopted birth dose HBV vaccination as part of the routine infant immunisation schedule [34]. Changes in the S protein can result in vaccine escape mutants (VEMs) [16,18], and also diagnostic escape mutations which result in false negative HBsAg testing [16]. Mutations in HBV Pol can also lead to amino acid changes in the Surface (S) protein due to overlapping reading frames (ORFs) in the genome [16]. Whilst the S protein mutation sG145R has been identified as the major VEM, recently other mutations in S protein have been associated with immune escape [16] Fig 1. There are very few data for VEMs in Africa, but in other settings of high endemicity, VEMs can be common, as evidenced by a reported prevalence of 28% in vaccinated HBV-infected children in Taiwan [35].

To date, no systematic review has assessed the geography and prevalence of HBV RAMs and VEMs in Africa. An understanding of the extent to which these mutations circulate in Africa is essential to improving HBV therapy in patients with and without HIV coinfection. We therefore set out to describe the frequency, co-occurrence and distribution of RAMs and VEMs in Africa, and to suggest whether changes are needed in recommendations for laboratory diagnostics and/or approaches to drug therapy or vaccine deployment. This will underpin further research to identify and track relevant mutations in these populations.

## METHODS

### Search strategy

Between October 2017 and January 2018, we searched the published literature, in MEDLINE (PubMed; https://www.ncbi.nlm.nih.gov/pubmed), SCOPUS (https://www.elsevier.com/solutions/scopus) and EMBASE (https://www.elsevier.com/en-gb/solutions/embase-biomedical-research). Our search strategy is detailed in S1 Table (documenting use of PRISMA criteria and selection of studies) and S2 Table (listing our search criteria). The earliest paper we identified on HBV drug resistance in Africa was published in 2007. We reviewed the titles and abstracts matching the search terms and only included those relating to drug or vaccine resistance in HBV infection, including only those that presented original data and had undergone peer review. All retrieved articles were in English, therefore no exclusion in relation to language was required.

For each publication we recorded reference, publication year, study design, sample size, study population, proportion of participants who tested HBsAg+ or HBV DNA+, country, year(s) of specimen collection, genotype identified, antiviral treatment, sequencing method, gene sequenced, number of sequenced samples, participant recruitment site and sequence accession number. Data were curated using MS Excel software (Microsoft, Redmond, WA).

### RAMs reported in published sequences not represented in primary studies

We expanded our search for evidence of RAMs by identifying publicly available HBV sequences from Africa, that had not been included in the results of our primary literature search. We used both the Hepatitis B Virus database (https://hbvdb.ibcp.fr/ [36] and Hepatitis Virus Diversity Research Alignments database (http://hvdr.bioinf.wits.ac.za/alignments/) [37].

### Analysis

In order to determine the prevalence of RAMs and VEMs, we first reported these using the denominator (total number of HBV positive patients) and numerator (total number of HBV positive patients with the specified mutation) as reported in published studies. We also pooled data by country in order to provide regional estimates. Downloaded sequences were managed using Sequence editor, database and analysis platform, SSE version 1.3, for analysis [38].

### Data visualisation

We developed an R package, gene.alignment.tables, for the visualisation of the sequence data in this study; this is available on Github [39] and can be used for visualising generic gene sequence datasets. The package was developed by University of Oxford’s Interactive Data Network and a specific instance of the visualisation is hosted as a Shiny app which can be viewed here: https://livedataoxford.shinyapps.io/1510659619-3Xkoe2NKkKJ7Drg/ [40].

## RESULTS

The initial search yielded 56 articles in MEDLINE, 150 in SCOPUS and 150 in EMBASE. Of these, 32, 136 and 119 were excluded from search results of MEDLINE, SCOPUS and EMBASE respectively, as they did not did not meet the inclusion criteria. After de-duplication, 37 articles were included. 27 articles identified from MEDLINE, SCOPUS and EMBASE were identical; five unique articles were included from EMBASE, four from SCOPUS and one from MEDLINE. A total of 37 articles were downloaded in full (S1 Table (part II); S3 Table).

### Study characteristics

Epidemiological data for HBV represented by the 37 studies we identified are summarised in Table 1. Studies included were from Southern Africa (Botswana, Mozambique, South Africa, Zambia and Zimbabwe), East Africa (Ethiopia, Kenya, Malawi, Sudan and Uganda), West Africa (Cote d’Ivoire, Gambia, Ghana, Guinea-Bissau and Nigeria) and Central Africa (Cameroon, Gabon). There was considerable heterogeneity in recruitment protocols and exposure to anti-viral treatment. Twenty-six studies recruited from hospitals, three studies recruited from the community [8,41,42] and eight studies did not specify where recruitment was undertaken [10,43–49]. All studies were observational.

**Table 1:** Prevalence of HBsAg and HBeAg from 37 studies of HBV drug resistance in Africa

|  | Study (Reference) | Country | Characteristics of study population | HIV co-infection status of cohort<sup>a</sup> | HBV prevalence in this cohort (reported as HBsAg prevalence unless otherwise specified) | HBeAg prevalence among HBV-positive individuals |
| --- | --- | --- | --- | --- | --- | --- |
| **East Africa** | Deressa et al 2017 [50] | Ethiopia | Patients attending outpatient ART clinic at tertiary referral university hospital | + | 17/308 (6%) | Not reported |
|  | Hundie et al 2016 [41] | Ethiopia | Stored plasma samples from HBV infected blood donors obtained from blood bank centres | ± | 391/391 (100%) | Not reported |
|  | Day et al 2013 [8] | Kenya | Longitudinal cohort study of female sex workers in an urban setting | + | 11/159 (7%) | 6/11 (55%) |
|  | Kim et al 2011 [9] | Kenya | Individuals from an urban centre enrolled in randomised controlled trial of adherence to ART | + | 27/389 (7%) | 24/27 (89%) |
|  | Mabeya et al 2017 [13] | Kenya | Individuals seeking treatment at the comprehensive HIV Clinic at tertiary referral university hospital | + | 29/400 (7%) | Not reported |
|  | Auodjane et al 2014 [20] | Malawi | Individuals starting ART treatment at a tertiary referral university hospital | + | 133/1117 (12%) | 67/133 (50%) |
|  | Galluzzo et al 2012 [45] | Malawi | Pregnant women enrolled in a PMTCT study on safety and pharmacokinetics of antiretroviral drugs | + | 21/21 (100%) | 7/21 (33%) |
|  | Mahgoub et al 2011 [42] | Sudan | Plasma samples from blood donors from capital city in Sudan | ± | 16/404 (4%) | Not reported |
|  | Yousif et al 2014 [47] | Sudan | Individuals seeking treatment at a AIDS care unit and HIV treatment centre | + | 96/358 (27%)<sup>b</sup> | 32/ 50 (64%) |
|  | Calisti et al 2015 [51] | Uganda | All HIV patients attending a regional referral hospital | + | 109/2820 (4%) | Not reported |
| **West Africa** | Boyd et al 2015 [52] | Cote d'Ivoire | Individuals enrolled in randomised multi centre trials of benefits and risks of early ART initiation | + | 259/ 2465 (11%) | 39/168 (23%) |
|  | Archampong et al 2017 [12] | Ghana | Serum samples from HBV-HIV co-infected patients collected at tertiary referral university hospital | + | 235/235 (100%) | Not reported |
|  | Chadwick et al 2012 [53]<sup>d</sup> | Ghana | Stored sera from all adult patients attending the HIV clinic at a tertiary referral university hospital | + | 143/371 (39%) | Not reported |
|  | Geretti et al 2010 [54]<sup>d</sup> | Ghana | Consecutive serum samples collected from unselected HIV-infected patients attending a tertiary referral university hospital | + | 140/838 (17%) | 37/140 (26%) |
|  | Ndow et al 2017 [55] | Gambia | Individuals attending HIV clinic | + | 23/187 (12%) | Not reported |
|  | Stewart et al 2011 (44) | Gambia | Individuals receiving HAART; recruitment site not specified | + | 70/ 570 (12%) | 6/21 (29%) |
|  | Langhoff Hongo et al 2014 [56] | Guinea Bissau | Patients attending outpatient ART clinic at tertiary referral university hospital | + | 94/576 (16%) | 16/94 (17%), HDV prevalence : 18/72 (25%) |
|  | Faleye et al 2015 [57] | Nigeria | Pregnant women attending antenatal clinics from two tertiary university hospitals | ± | 15/272 (6%) | Not reported |
| **Central Africa** | Gachara et al 2017 [58] | Cameroon | Patients attending outpatient ART health centre | + | 20/337 (6%)<sup>c</sup> | Not reported |
|  | Kouanfack et al 2012 [59] | Cameroon | Patients attending outpatient ART clinic at tertiary hospitals | + | 54/552 (10%) | Not reported |
|  | Magoro et al 2016 [60] | Cameroon | Patients attending outpatient ART health centre | + | 116/445 (26%) | 16/102 (16%) |
|  | Bivigou-Mboumba et al 2016 [61] | Gabon | Patients attending outpatient ART clinic | + | 71/762 (9%) | Not reported |
|  | Bivigou-Mboumba et al 2018 [62] | Gabon | Patients attending HIV care centers | + | 43/487 (9%) | Not reported |
| **Southern Africa** | Anderson et al 2015 [48] | Botswana | Stored plasma samples of HIV/HBV co-infected individuals collected from studies conducted in a Research Institution | + | 81/81 (100%) | Not reported |
|  | Matthews et al 2015 [63] | Botswana | Women attending antenatal and paediatric clinics | ± | 17/443 (4%) | 16/60 (27%); HDV: negative |
|  | Chambal et al 2017 [64] | Mozambique | Patients attending outpatient ART health centre | + | 47/518 (9%) | Not reported |
|  | Wandeler et al 2016 [14] | Mozambique | Individuals starting ART treatment at urban clinic in Mozambique and rural clinic in Zambia | + | 78/1032 (8%) | 24/168 (14%) |
|  | Andersson et al 2013 [65] | South Africa | Stored serum of women infected with HIV enrolled in an Antenatal Sentinel HIV and Syphilis Prevalence Survey | ± | 97/3089 (3%) | 17/94 (18%); HDV: negative |
|  | Amponsah-Dacosta et al 2015 [43] | South Africa | Stored serum of individuals exposed to HBV participating in a health facility-based hepatitis B serosurvey conducted at a provincial level. | ± | 33/201 (16%) | Not reported |
|  | Amponsah-Dacosta et al 2016 [49] | South Africa | Individuals due to HAART initiation enrolled in longitudinal study | + | 5/5 (100%) | 5/5 (100%) |
|  | Hamers et al 2013 [10] | South Africa | Individuals enrolled in a multicentre prospective study of ART resistance monitoring | + | 37/175 (21%) | Not reported |
|  | Gededzha et al 2016 [66] | South Africa | Stored sera from HBV infected individuals attending a tertiary referral university hospital | ± | 8/9 (89%) | Not reported |
|  | Makondo et al 2012 [46] | South Africa | Stored sera from HIV infected individuals prior to ART initiation, recruitment site not specified | + | 71/298 (24%)<sup>b</sup> | Not reported |
|  | Matthews et al 2015 [63] | South Africa | Women attending antenatal and paediatric clinics in South Africa and Botswana | ± | 49/507 (10%) | Not reported; HDV: negative |
|  | Powell et al 2015 [67] | South Africa | Stored serum samples of individuals infected with HIV receiving care at a tertiary university hospital | + | 37/394 (9%) | Not reported |
|  | Selabe et al 2007 [68] | South Africa | Individuals infected with HBV admitted at tertiary University hospital | ± | 35/35 (100%) | Not reported |
|  | Selabe et al 2009 [69] | South Africa | Individuals infected with HBV admitted at tertiary University hospital | - | 17/17 (100) | 9/17 (53%) |
|  | Wandeler et al 2016 [14] | Zambia | Individuals starting ART treatment at urban clinic in Mozambique and rural clinic in Zambia | + | 90/797 (11%) | 24/168 (14%) |
|  | Hamers et al 2013 [10] | Zambia | Individuals enrolled in a multicentre prospective study of ART resistance monitoring | + | 55/523 (11%) | Not reported |
|  | Baudi et al 2017 [70] | Zimbabwe | Stored plasma samples of individuals attending HIV support clinic | + | 19/176 (11%) | Not reported |

HBsAg and HBeAg prevalence were determined from 37 studies (Treatment naïve: n= 8 studies, 566 individuals with HBsAg; Treatment experienced: n= 19 studies, 1243 individuals with HBsAg; Mixed regimen where some were treatment experienced, naïve or treatment status not specified: n= 10 studies, 1046 individuals with HBsAg). Studies were identified by a systematic literature search of HBV resistance associated mutations (RAMs) and vaccine escape mutations (VEMs) from African cohorts published between 2007 and 2017 (inclusive).
<sup>a</sup> HIV status is designated '+' whole cohort HIV-positive; '±' some of cohort HIV-positive; '-' none of cohort HIV-positive
<sup>b</sup> HBV prevalence in these cohorts was reported using HBV DNA detection rather than HBsAg
<sup>c</sup> Occult HBV prevalence reported in these cohorts
<sup>d</sup> These two studies recruited from the same overall cohort in Ghana.

Study populations were categorised as follows:

- HBV/HIV coinfected patients: (n=28 studies), [8–10,12–14,20,43–48,50–56,58–, –62,64,67,70];
- HBV infected with and without HIV coinfection: (n=8 studies), [41–43,57,63,65,66,68];
- Chronic HBV monoinfection: (n=1 study), [69].

Antiviral treatment exposure varied as follows:

- Treatment-naïve: (n=8 studies), [14,46–48,63,64,68,70];
- 3TC-based regimen only: (n=10 studies), [8,9,20,44,45,49,53,59,61,69];
- Regimens including 3TC or TDF: (n=6 studies), [10,13,51,52,55,65];
- Mixed regimen where some received 3TC, others TDF, while others left untreated; (n=7 studies), [12,50,54,56,60,62,66];
- Treatment regimen not specified: (n=6 studies), [41–43,57,58,67].

HBV amino acid polymorphisms were studied from within the following proteins;

- Pol only (n=13 studies), [8,9,12–14,20,44,53,55,56,59,63,68]; only one of these used a deep sequencing method [20];
- S only (n=3 studies), [43,57,65];
- Pol and S (n=12 studies),[10,41,45,48,51,52,54,58,60,62,64,67];
- Pol, S and PC/BCP (n=4 studies), [47,50,61,69];
- S and PC/BCP region (n=3 studies), [42,46,70];
- Whole genome (n=2 studies), [49,66].

All studies, except for two [53,68], specified the HBV genotype (S3 Table & S2 Fig).

### Prevalence of HBsAg, HBeAg and HDV coinfection

The prevalence rates of HBsAg in these study cohorts ranged from 3%-26%; however, the populations included were highly selected and therefore not necessarily representative of the general population, particularly as a result of a strong bias towards HIV-infection (Table 1). Only three studies included in this review reported on HDV prevalence: two studies did not detect any HDV antibodies [63,65], whereas the other study reported a HDV prevalence of 25% in Guinea-Bissau [56].

### RAMs identified in African cohorts

The co-occurrence and distribution of HBV RAMs and VEMs are summarised according to the region where they were identified (Fig 2). This illustrates the patchy and limited data that are available, with South Africa, Ghana and Cameroon best represented, but with large areas (especially in northern and central Africa) not represented at all in the literature.

**Figure 2:**
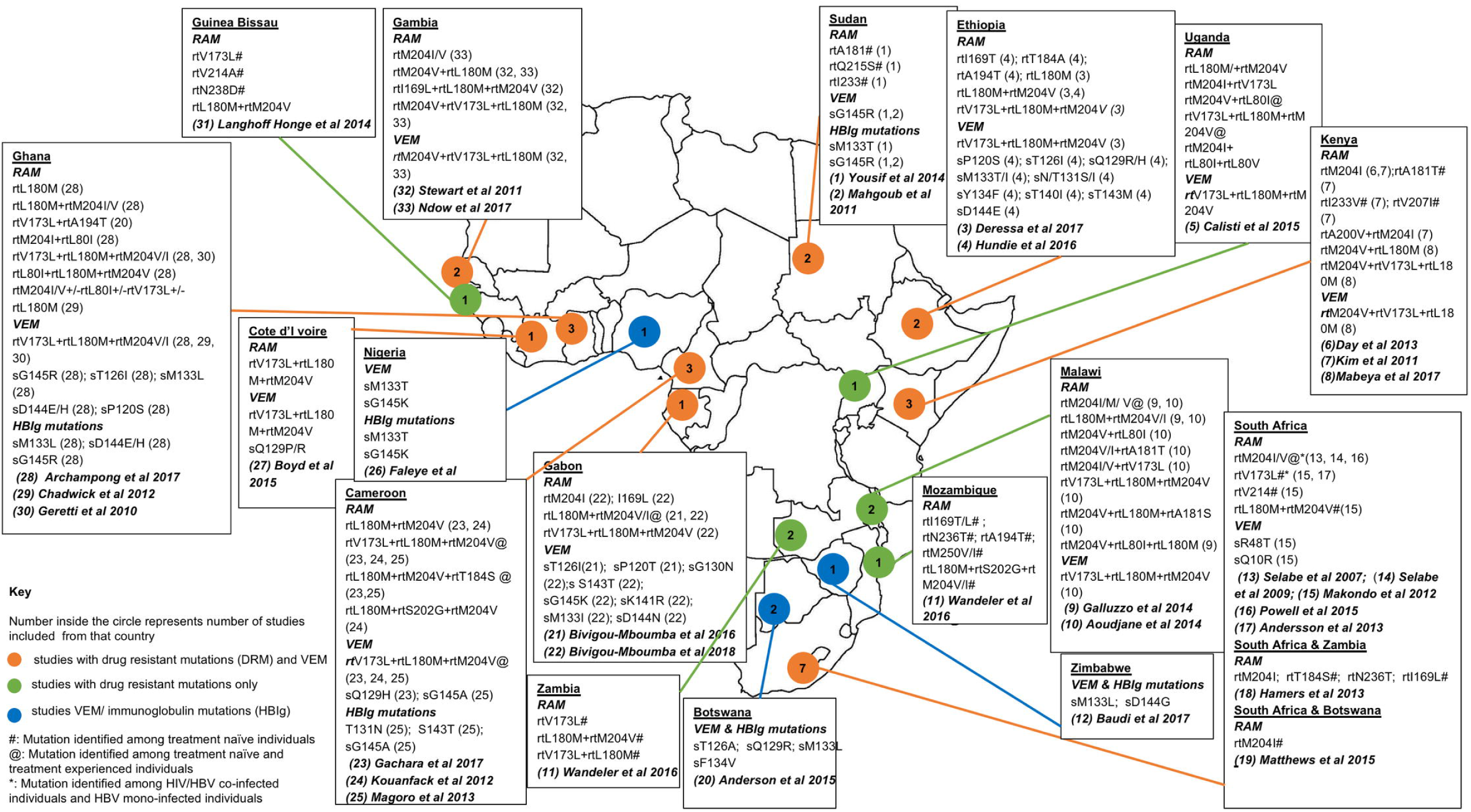
Annotated map to summarise HBV drug Resistance Associated Mutations (RAMs) and Vaccine Escape Mutations (VEMs). Mutations identified from 33 studies of African cohorts published between 2007 and 2017 (inclusive). Four studies identified by our systematic literature review were not represented here as they did not report any RAMs.

Although 35 studies specified the HBV genotype, it was only possible to group RAMs according to genotype in fourteen studies [8,9, 13, 14,44,46,47,50,51,56,60–62,69] (S1 Fig; S2 Fig). The remaining 21 studies generally reported the genotypes identified, but did not specifically state the genotype of HBV within which RAMs were identified.

We have developed an interactive tool to display the genomic positions of RAMs identified through our literature review alongside relevant metadata. This can be accessed on-line here: https://livedataoxford.shinyapps.io/1510659619-Xkoe2NKkKJ7Drg/ [40].

Overall, the most prevalent RAM was rtM204V/I in both treatment experienced and treatment naïve individuals, and occurring either alone or in combination with other polymorphisms rtL80I/V, rtV173L, rtL180M, rtA181S, rtT184S, rtA200V and/or rtS202S (Fig 3); mutations among individuals with and without exposure to HBV therapy are listed in S4 Table and S5 Table, respectively). This mutation was present in 29 studies at a highly variable prevalence of between 0.4% [12] and 76% [69]. Across all cohorts, the mutation was present in 208/2569 (8%) of all individuals represented. The mutation, by itself, was most prevalent in South Africa; on pooling data for three studies from this setting, it was present in both treatment experienced and treatment naïve patients (n=13/17, 76% [69] and n=16/72, 22% [67,68] respectively). In addition to South Africa, rtM204I/V was also frequent in Malawi among treatment experienced patients (n=24/154, 16% [20,45]) (Fig 3), and in genotype non-A infection: in this setting, the mutation was detected in genotype C infection (n=2/17, 12% [69]) (S2 Fig).

**Figure 3:**
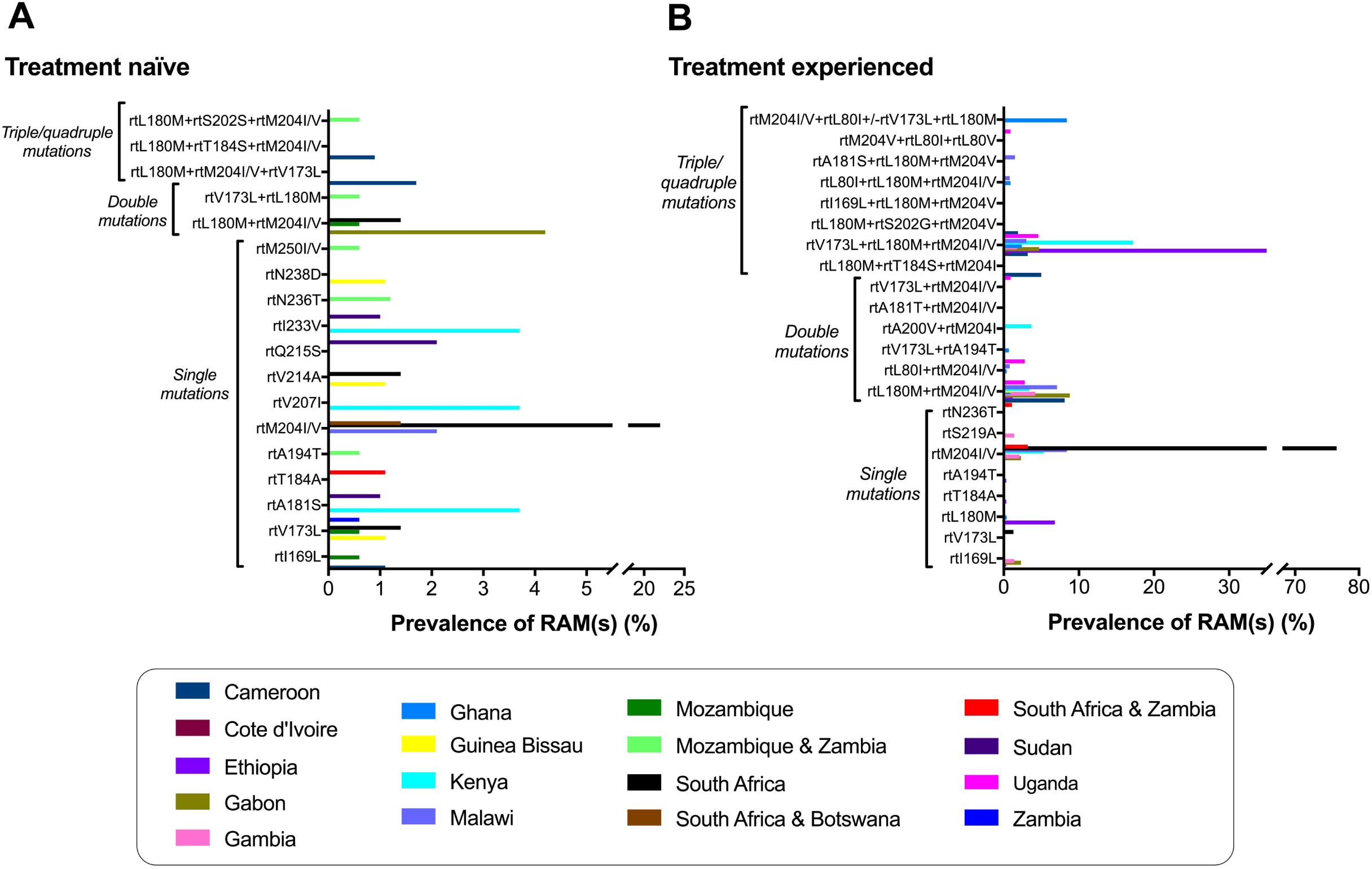
Prevalence of HBV resistance associated mutations (RAMs) in Pol/RT proteins among HBV infected patients in Africa. These data are derived from 27 studies of HBV drug resistance in Africa published between 2007 and 2017 (inclusive). The countries represented are listed in alphabetical order. A detailed summary of RAMs identified from each study is presented (Fig 2, S4 Table, S5 Table). Prevalence of RAMs for a specific country was determined by grouping all studies from that country that reported a specific mutation. We used all individuals who tested HBsAg positive to generate a denominator in order to provide a conservative estimate of RAM prevalence, and the numerator was the total number of individuals with that specific mutation from these studies. A: treatment naïve; B: treatment experienced.

The rtM204I/V mutation by itself confers resistance to 3TC; in combination with A194T it may also be associated with reduced efficacy to TDF, and in combination with L180M and V173L with vaccine escape, through corresponding substitutions in the surface antigen sites targeted by neutralising antibodies. Although TDF has a high genetic barrier to resistance, and is associated with reliable suppression of HBV viraemia [7,10,21,22], mutations rtN236T and rtA194T, which have been linked with resistance to both TDF and ADV [7], have been identified in Southern Africa in both treatment naïve [14] and treatment experienced [10] patients.

WHO guidelines recommend a first-line regimen including TDF in HIV/HBV coinfected patients [6], and the South African Department of Health HIV/AIDS treatment guideline included TDF as first-line regimen from 2010 [71], however we found a minority of studies (9/37, 24%) reporting TDF-containing regimens for HIV/HBV coinfected individuals. As anticipated, most of the studies that did use TDF were carried out after 2010, whereas those that used 3TC were generally earlier (S3 Table).

From this dataset, it is difficult to ascertain whether RAMs are genuinely more prevalent in genotype A infection, or this simply reflects enrichment of genotype A in sub-Saharan African populations (S2 Fig). Interpreting RAMs according to sub-genotypes was difficult since most studies did not specify sub-genotype and others did not indicate which RAMs were identified in which genotype. Of concern is the detection of RAMs even in reportedly treatment naïve individuals (Fig 3 & S4 Table), suggesting that RAMs are being transmitted. A study in South Africa that recruited 3TC-naïve HBV infected adults with or without HIV, reported rtM204I in 13/35 (37%) individuals [68].

### HBV RAMs in published sequences from Africa

We searched the Hepatitis B Virus database and GenBank to identify HBV sequences derived from Africa, from studies not already included in our review. We identified an additional 69 isolates: 23 had undergone full length genome sequencing whereas 46 isolates represented either the polymerase (n=3) or S region (n= 43) of the HBV genome Table 2. To avoid duplication of results, we excluded fourteen studies already identified by our literature review that had submitted their sequences to GenBank (S3 Table). RAMs in the additional 69 isolates were as follows:

- rtM204V in genotype A (2/69, 2.9% of sequences), this occurred in combination with rtL180M;
- rtM204V + rtL180M in genotype E (1/69, 1.5%);
- rt180M + rtA181V in genotype E (1/69 (1.5%);
- rtQ215S identified in genotype D (4/69, 5.8%).

All these mutations are associated with 3TC resistance; rtA181V has also been associated with reduced susceptibility to TDF [7,15].

In the S gene, the most prevalent mutations were:

- sD144A/E/G occurring in genotype A (6/69, 8.7%), D (10/69, 14.5%) and E (7/69, 10.1%) associated with VEM;
- sI110L occurring in genotype A (3/69, 4.3%), D (4/69, 5.8%) and E (11/69, 15.9%) associated with immunoglobulin resistance.

**Table 2:**
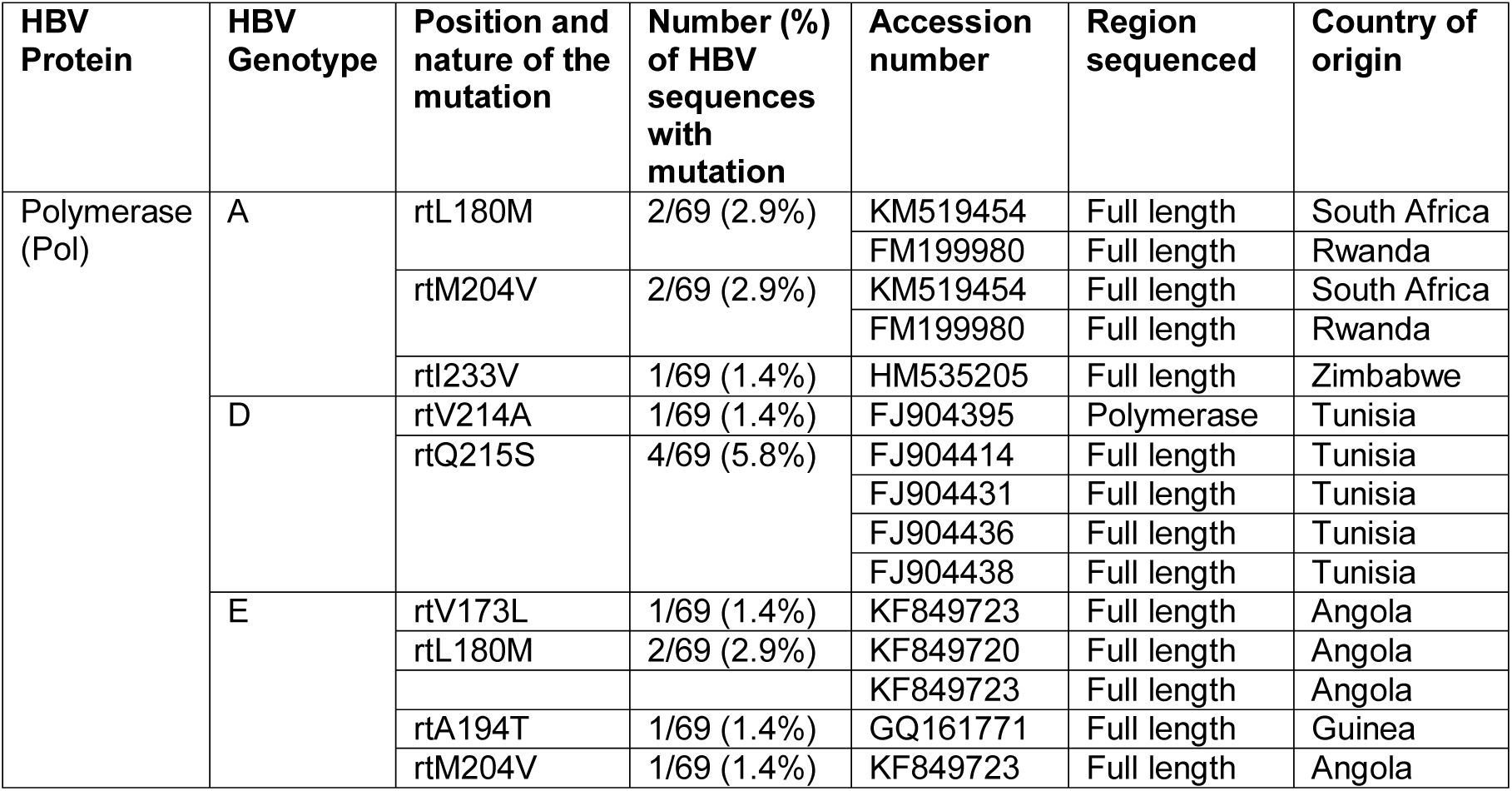
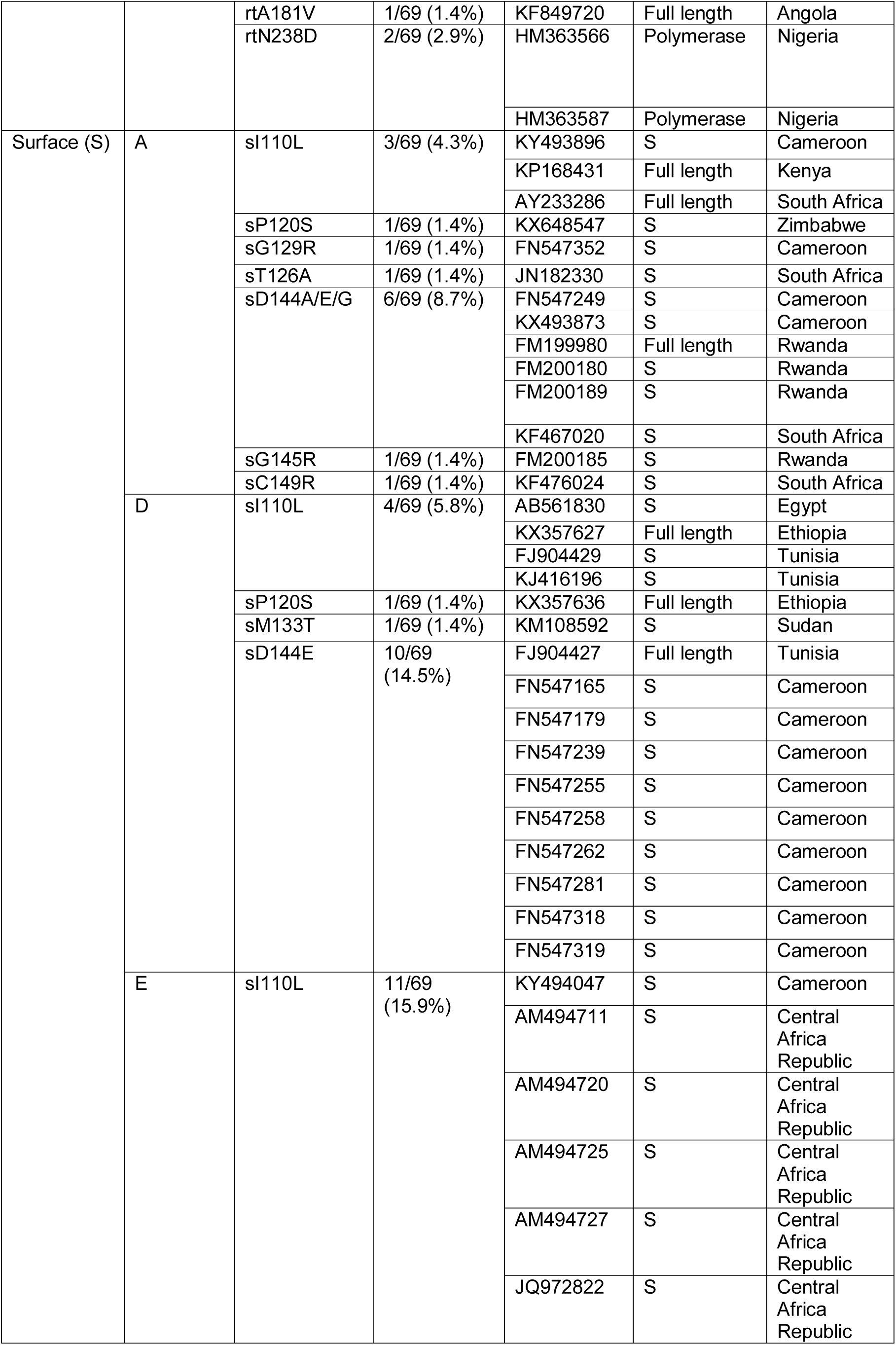
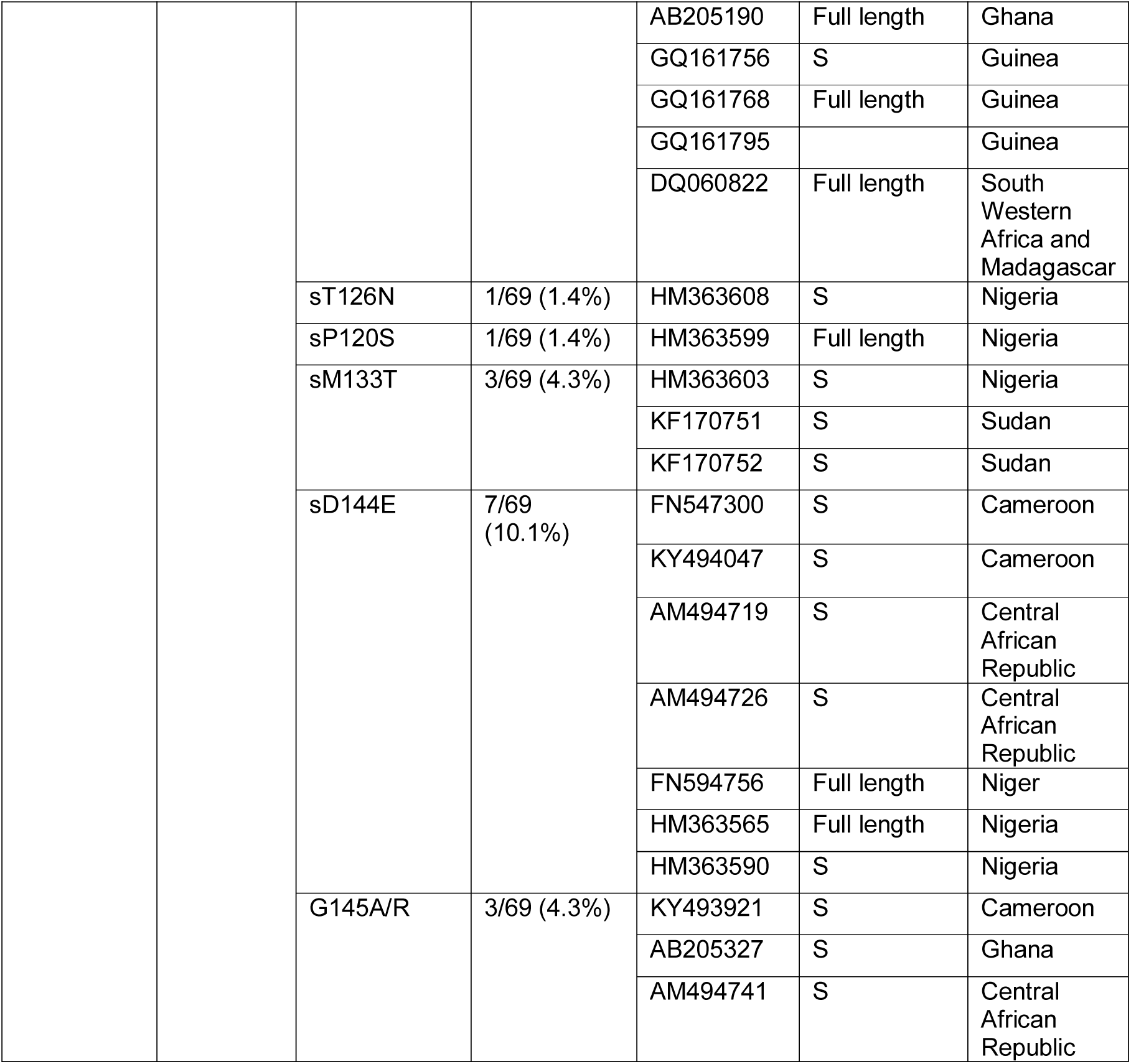
HBV drug resistant mutations (RAMs) identified from HBV genome sequences from Africa downloaded from the Hepatitis B Virus database (https://hbvdb.ibcp.fr/) [36] and GenBank database (http://hvdr.bioinf.wits.ac.za/alignments/) [37]

### VEMs

VEMs were identified in Central, East, West and Southern Africa (Fig 2). However, it was not possible to ascertain whether individuals harbouring these mutations had been vaccinated against HBV infection. The most common VEM was the triple mutation rtV173L + rtL180M + rtM204I/V, found in the *Pol* gene. This suite of mutations was identified in 14 studies [12,13,20,44,50–55,58–60,62], at a pooled prevalence of 4% (57/1462). Another significant VEM, sG145K/R [16], was identified in six studies [12,42,47,57,60,62] and sM133L/T, associated with VEM, immunoglobulin and diagnostic escape mutation [12,48], was identified in seven studies [12,41,47,48,57,62,70] (Fig. 2).

## DISCUSSION

### Summary

To our knowledge, this is the first systematic review that assesses RAMs and VEMs for HBV in Africa. The high rates of HBV infection among HIV infected individuals in some locations including Cameroon [60] and South Africa [10] could be an indication that HBV infection has been previously under-reported, possibly due to lack of routine screening, poor awareness, stigma, high costs and limited clinical and laboratory infrastructure [4,8–10,45,53]. The literature suggests a widespread exposure of the HIV-infected population to 3TC-based treatment. This may be changing over time in line with current ART treatment recommendations (regimens for Africa summarised in S6 table), but the introduction of TDF-based regimens for HIV treatment has been inconsistent, and TDF monotherapy is not consistently available for HBV infection in the absence of HIV.

In keeping with other settings, the most common RAM identified here was rtM204I/V, either alone or in combination with compensatory mutations rtL180M ± rtV173L. Of concern, rtM204V/I was seen in 76% of treatment experienced patients [69] and 22% of treatment naïve patients [67,68] in South Africa. A review of worldwide incidence of RAMs among treatment naïve patients also described rtM204V/I as the most frequent, but with a much lower prevalence of 5% [72]. The contribution of unreported or undocumented 3TC exposure in the reportedly treatment naïve populations remains to be determined. A European study demonstrated that the most frequent primary mutation was rtM204V/I, found in 49% of treatment experienced patients [73], while in China rtM204I, rtN236T and rtL180M+rtM204V+rtV173L/rtS202G were also the most prevalent RAMs [74].

The triple mutation rtM204V + rtL180M + rtV173L has been identified in East, West and Central Africa [20,44,51–54,59]. This combination of polymorphisms is associated with both vaccine escape and resistance to 3TC and other-nucleoside analogues [20,44,51,54,59,60]. Interestingly, this triple mutation has not been reported in the Southern African region to date, which is likely to reflect the composition of the study populations.

### Clinical and public health significance of RAMs

Apart from the nature of drug being used for HBV treatment, other predictors of HBV drug resistance include HBV viral load, HBV intra host heterogeneity, HBeAg status, host body mass index and serum alanine aminotransferase (ALT) activity [20,75,76]. Individuals with rtM204V/I plus compensatory mutations typically exhibit high HBV DNA levels [20] and are therefore highly infectious to others. The spread of RAMs may lead to a rise in drug resistance in treatment naïve chronic HBV infection, representing a substantial challenge for Africa and highlighting an imperative to ensure routine use of TDF in preference to, or in combination with, 3TC-based therapy.

Although these data provide a preliminary picture of the prevalence of RAMs in some settings, there are no recommendations to stipulate any specific prevalence threshold above which HBV drug resistance mutations represent a significant barrier to successful treatment at a population level, and/or RAM prevalence thresholds that should trigger a switch to alternative first-line therapy. For HIV, surveillance for transmission of RAMs is based on screening recently infected, treatment naive individuals, and classifies drug resistance using thresholds of <5%, 5-15%, and >15% to stratify the risk to public health [77]. Similar thresholds and recommendations for HBV could help to underpin the assimilation of epidemiological data and to unify treatment approaches.

### TDF resistance

The identification of mutations associated with reduced TDF susceptibility are of concern, as they suggest the potential for increasing prevalence of polymorphisms that confer partial or complete viral escape from a drug that to date has not been widely associated with resistance. There is now potential for increasing selection of TDF resistance as this drug becomes more widely used. However, as a new first line single tablet option incorporating 3TC, TDF and Dolutegravir (DTG) (triple therapy abbreviated to ‘LTD’) emerges as a recommended option for HIV treatment in Africa, surveillance is needed to determine the clinical outcomes for HBV [78].

If clinically significant, TDF resistance mutations may still represent a particular problem for many African settings, as resource constraints make it unrealistic to provide baseline screening for RAMs, or to monitor patients on treatment with serial viral load measurements. Despite these potential concerns, it has been shown that TDF is effective even in the presence of RAMs and that there is comparable efficacy among 3TC-experienced and NA-naïve patients [79].

### VEM

VEM were identified in 16 different countries in East, West, Central and Southern Africa. Information on vaccine exposure was not available, but there are two strands of evidence to support significant population exposure to HBV vaccination. First, vaccination has been progressively rolled out in most countries in sSA since the mid-1990’s; second, most HBsAg mutations reported by these studies are located within the common immunodominant B cell epitope (aa 124-147) in which selection of polymorphisms is associated with HBV vaccination [80,81].

VEM have been more robustly reported from Asia, in settings where the HBV infant vaccination programme is well established; for example, in Taiwan, VEM prevalence among vaccinated children increased from 7.8% to 23.1% within 15 years of the launch of the universal vaccine program, although the decline in VEM prevalence thereafter may be partly related to a smaller HBV carrier pool [80]. HBV infection despite immunoprophylaxis can occur either as a consequence of MTCT of pre-existing VEM, or as a result of *de novo* selection of escape mutations from vaccine-induced immune responses, particularly in the setting of delayed vaccination [80,81].The HBV genotype sequence used for vaccines may potentially have an influence on immunogenicity against non-vaccine genotypes, but there are limited data to support this [82]. Only 11 African countries recommend the first HBV vaccine dose at birth, in contrast to the majority of African countries in which HBV vaccination is delayed until 6 weeks of age [33]. It is likely that this delay not only provides a window of infection but also increases the possibility of transmitted VEM and/or emergence of new escape mutations.

High maternal HBV viral load and immunosuppression are other risk factors associated with VEM among infants [80]; both of these are pertinent for emergence of VEMs in Africa given that HBV viral load testing is not routinely available, and HIV is highly prevalent in some populations. Effective PMTCT strategies in Africa, including screening and treating antenatal women, increasing access to viral load monitoring, and introducing HBV birth dose vaccine will help to decrease the prevalence of VEM [4,33,83].

### HDV/HBV coinfection

One study from our literature review reported a high HDV prevalence of 25%; however, in this cohort, RAMs occurred in individuals with HBV monoinfection [56]. Given that HDV is characteristically associated with decreased HBV replication [84], it is possible that emergence of HBV RAMs is altered in this setting. However, as the true prevalence and impact of HDV in sSA is not known [85], further studies are needed to determine the impact of HDV coinfection on HBV RAMs.

### Limitations of current data

Screening for HBV infection is not routinely performed in many African settings and therefore the true prevalence and characteristics of HBV infection are not known [4,7–10]. We identified very few published studies; only a minority of patients had HBV sequencing undertaken, and there were no data from certain regions of Africa. This highlights the substantial problem of HBV neglect in Africa, and a specific blind-spot relating to sequence data [4]. Identifying the true prevalence of resistance mutations, and characterising the populations in which these are selected and enriched, is currently not possible due to sparse data and lack of clear descriptions of the denominator population. Most such studies do not perform a truly systematic assessment, but focus on high risk groups – particularly including those with HIV/HBV coinfection: of the 37 studies included here, only one exclusively reported on participants who were HBV mono-infected [69]. Although we have made every effort to assimilate the relevant data to build up a regional picture for Africa, the heterogeneity between studies makes it difficult to draw robust conclusions from pooled data. These findings are a reflection of the little attention paid towards the burden of this disease in Africa and the neglect in robust epidemiological data.

Only nine studies undertook a longitudinal approach to detection of drug resistance [8–10,20,44,45,49,52,53]. The results of the other 28 studies that undertook a cross-sectional approach could be skewed by the timing of recruitment of study participants, with a risk of under-representation of drug resistance if screening is undertaken only at baseline, and potentially an over-representation if screening is undertaken in patients with HIV coinfection, who are more at risk of advanced disease and prolonged drug exposure. As most of these studies recruited individuals from hospital settings, this raises the latter possibility.

Mutations across the whole genome might be relevant in determining resistance [86]. However, most of the included studies analysed only defined genes from within the HBV genome; only two sequenced the whole genome, and these determined consensus sequence. This potentially results in an under-representation of RAMs and VEMs that may be present as low numbers of quasispecies, but could become significant if selected out by exposure to drug or vaccine.

In studies that reported RAMs among treatment naïve individuals, the literature suggests that sequence analysis was performed prior to ART initiation. However, we cannot exclude the possibility that that some of these participants had prior ART exposure. Due to the nature of the cohorts that have been studied, most of the RAMs identified were from HIV/HBV coinfected individuals. It is possible that HIV increases the risk of HBV RAMs both in terms of drug exposure, and also as a function of increased HBV viral loads. A study from Malawi demonstrated the rapid emergence of 3TC resistance in HIV coinfection, with virtually all treatment naïve HBeAg positive individuals starting antiviral treatment showing emergence of rtM204I by six months. Likewise, a study carried out in Italy revealed that patients with HIV coinfection were more likely to harbour the rtM204V mutation and to show multiple mutations compared to HBV monoinfected patients [87]. It would be worth further exploration of this observation in Africa, as there are currently very limited data.

### Challenges and opportunities for Africa

A major challenge for Africa is to improve coverage rates of infant vaccination, deploy catch-up vaccination programmes for older children and adults, adopt widespread screening and develop treatment programmes for HBV. While HBV vaccine is effective, gaps in vaccine coverage in Africa can be demonstrated by the high perinatal transmission rate of HBV in sSA (estimated at 38% among women with a high HBV viral load) and the observation that up to 1% of newborns in sSA are still infected with HBV [88]. Sustained efforts are required to build robust PMTCT programmes that deliver screening and treatment for antenatal women, and timely administration of HBV birth vaccine for their babies [33,83].

Although the WHO recommends monitoring for the development of drug resistance once on therapy [6], implementation remains challenging as viral load monitoring and sequencing are both rarely available [7]; despite the advancement and availability of HIV testing and monitoring, in many settings it remains uncommon to monitor HIV viral load after ART initiation [89,90]. Affordable, accessible and sustainable platforms for quantifying both HIV and HBV viral loads remain an important priority for many settings in Africa, given the lack of on-treatment monitoring in many settings. Given the simplicity and relative ease of collection, preparation and transport of dried-blood-spot (DBS) samples [91], adopting DBS testing could improve access to HBV diagnosis, viral load monitoring and linkage to care, especially in areas with limited access to laboratory facilities.

Development of a cheap, rapid test for the detection of the most frequently observed RAMs and VEMs should be considered as a potentially cost-effective strategy for Africa. Proof of principle for a rapid test for diagnosis and detection of resistance has been demonstrated by the GeneXpert MTB/RIF assay for *Mycobacterium tuberculosis* (MTB) [92]. A similar approach has been applied for HBV through use of a multiplex ligation-dependent probe real time PCR (MLP-RT-PCR) [93]. Although this assay is able to detect RAMs quickly and cheaply, there are still limitations as the test requires high viral load samples, is based on detection of known RAMs from within discrete regions of the genome, and may not identify RAMs that are present as minor quasispecies.

New metagenomic sequencing platforms, such as Illumina and Nanopore, provide the opportunity for whole deep genome sequencing, which can reveal the full landscape of HBV mutations in individual patients, quantify the prevalence of drug resistance mutations among HBV quasi-species, and determine the relationship between these polymorphisms and treatment outcomes [87]. Nanopore technology also has the potential to develop into an efficient point of care test that could detect viral infection and coinfection, as well as determining the presence of VEMs and RAMs [94], but is currently limited by cost and concerns about high error rates.

There have been few studies looking at the correlation between genotype, clinical outcomes of disease, response to antiviral therapy and RAMs/VEMs, but none from Africa. Studies outside Africa have shown that genotype A is more prone to immune/vaccine escape mutants, pre-S mutants associated with immune suppression, drug associated mutations and HCC in HIV/HBV coinfected participants [46,87,95]. Studies investigating the role of genotypes in predicting response to antiviral therapy and their association with various types of mutations are urgently needed in Africa, particularly in light of the high frequency of genotype A infection and high population exposure to antiviral agents that have been rolled out over the past two decades as a component of first-line ART.

Existing infrastructure for diagnosis, clinical monitoring and drug therapy for HIV represents an opportunity for linkage with HBV care. Particularly in settings of limited resource, joining up services for screening and management of blood-borne virus infection could be a cost-effective pathway to service improvements.

### Conclusions

This review highlights the very limited data for HBV RAMs and VEMs that are available from Africa. Scarce resources resulting in lack of diagnostic screening, inconsistent supply of HBV drugs and vaccines, and poor access to clinical monitoring contribute to drug and vaccine resistance, potentially amplifying the risk of ongoing transmission and adding to the long-term burden of HBV morbidity and mortality in Africa. We call for urgent action to gather and analyse better data, particularly representing the HBV monoinfected population, and for improved access to TDF.

HBV RAMs and VEMs have been identified in several African countries among HIV/HBV coinfected and HBV monoinfected patients, before and during treatment with NAs but the data are currently insufficient to allow us to form a clear picture of the prevalence, distribution or clinical significance of these mutations. Overall, the data we describe suggest a significantly higher prevalence of drug resistance in some African populations than has been described elsewhere, and that is not confined only to drug-exposed populations, highlighting an urgent need for better population screening, assessment of HBV infection before and during therapy, and increasing roll out of TDF in preference to 3TC. At present, TDF accessibility is largely confined to HIV/HBV coinfected individuals; we now need to advocate to make monotherapy available for HBV monoinfected individuals. However, there are uncertainties as to whether its long-term use might result in nephrotoxicity, and potentially in an increase in selection of TDF RAMs.

We should ideally aim for the goals of a combined HBV test that includes diagnosis of infection, genotype and presence of RAMs/VEMs; new sequencing platforms such as Nanopore make this technically possible, although cost remains a significant barrier at present. Sustainable long-term investment is required to expand consistent drug and vaccine supply, to provide screening infection and for drug resistance, and to provide appropriate targeted clinical monitoring for treated patients.

## SUPPORTING INFORMATION

**S1 Fig: HBV drug Resistance Associated Mutations (RAMs) grouped according to genotype.** Data summarised from fourteen studies published between 2009-2017 (inclusive). 21 studies were not represented here as they did not specifically indicate which genotype individuals with RAMs belonged to. **Available at** https://doi.org/10.6084/m9.figshare.5774091 [96].

**S2 Fig: Distribution of HBV genotypes and prevalence of HBV resistance associated mutations (RAMs) in Pol/RT proteins in geno-A and geno-non-A samples.** A: Distribution of HBV genotypes derived from 35 studies reporting resistance associated mutations (RAMs) in Africa published between 2009 to 2017 (inclusive); B: Prevalence of HBV resistance associated mutations (RAMs) in Pol/RT proteins in geno-A and geno-non-A samples. These data are derived from 14 studies of HBV drug resistance in Africa published between 2007 and 2017 (inclusive). 21 studies were not represented here as they did not specifically indicate which genotype individuals with RAMs belonged to. We had more geno-A samples represented than other samples, we therefore combined samples from other genotypes that had RAMs (B, C, D, E, D/E) to form geno-non-A samples. We then compared prevalence of Pol/RT mutation between geno-A samples to geno-non-A samples. Prevalence of RT/Pol mutations for a specific genotype(geno-A/geno-non-A) was determined by grouping all studies with geno-A/geno-non-A infection that reported a specific mutation; the denominator was the total number of individuals infected with geno-A/geno-non-A from these studies and the numerator was the total number of individuals infected with geno-A/geno-non-A with that specific mutation. **Available at** https://doi.org/10.6084/m9.figshare.5774091 [96].

**S1 Table: PRISMA (Preferred Reporting Items for Systematic Reviews and Meta-Analyses) criteria for a systematic review of hepatitis B virus (HBV) drug and vaccine escape mutations in Africa.** Available at https://doi.org/10.6084/m9.figshare.5774091 [96].

I. Checklist to demonstrate how PRISMA criteria (2009) have been met in this review;

II. Flow diagram illustrating identification and inclusion of studies for a systematic review of drug and vaccine resistance mutations in Africa.

**S2 Table: Details of search strategy used to identify studies on HBV resistance associated mutations (RAMs) and vaccine escape mutations (VEMs) conducted in Africa.** A: PubMed database; B: SCOPUS and EMBASE database. **Available at** https://doi.org/10.6084/m9.figshare.5774091 [96].

**S3 Table: Full details of 37 studies identified by a systematic literature search of HBV resistance associated mutations (RAMs) and vaccine escape mutations (VEMs) from African cohorts published between 2007 and 2017 (inclusive). Available at** https://doi.org/10.6084/m9.figshare.5774091 [96].

**S4 Table: HBV Pol/RT mutations among treatment-naïve HBV infected patients in Africa from 12 studies published between 2007 and 2017 (inclusive). Available at** https://doi.org/10.6084/m9.figshare.5774091 [96].

**S5 Table: HBV Pol/RT mutations among treatment-experienced HBV infected patients in Africa, from 25 studies published between 2009 and 2017 (inclusive). Available at** https://doi.org/10.6084/m9.figshare.5774091 [96].

**S6 Table: First line ART regimen for adults in Africa, and overlap with HBV therapy. Information derived from published ART guidelines in all cases where these are available in the public domain. This information was collated in May 2018. Available at** https://doi.org/10.6084/m9.figshare.5774091 [96].

